# Brain Transcriptional Profiles of Male Alternative Reproductive Tactics and Females in Bluegill Sunfish

**DOI:** 10.1101/025916

**Authors:** Charlyn G. Partridge, Matthew D. MacManes, Rosemary Knapp, Bryan D. Neff

## Abstract

Bluegill sunfish are one of the classic systems for studying male alternative reproductive tactics (ARTs) in teleost fishes. In this species, there are two distinct life histories: parental and cuckolder, encompassing three reproductive tactics, parental, satellite, and sneaker. The parental life history is fixed, whereas individuals who enter the cuckolder life history transition from the sneaker to the satellite tactic as they grow. For this study, we used RNAseq to characterize the brain transcriptome of the three male tactics and females during spawning to identify gene categories associated with each tactic and identify potential candidate genes influencing their different spawning behaviors. We found that sneaker males had higher levels of gene differentiation compared to the other two male tactics. Sneaker males also had high expression in ionotropic glutamate receptor genes, specifically AMPA receptors, which may be important for increased working spatial memory while attempting to cuckold parental males at their nests. Larger differences in gene expression also occurred among male tactics than between males and females. We found significant expression differences in several candidate genes that were previously identified in other species with ARTs and suggest a previously undescribed role for cAMP-responsive element modulator (*crem*) in influencing parental male behaviors during spawning.

## Introduction

Understanding the genes that influence variation in behavior can provide insight into how different behavioral phenotypes within populations evolve and are maintained. One important area of research on behavioral phenotypes focuses on alternative reproductive tactics (ARTs), which are found in a wide array of taxa [1–5]. ARTs typically consist of larger males practicing a “territorial” tactic that maintain and protect breeding territories and smaller “sneaking” males that sneak fertilizations rather than compete with territorial males [6]. The mechanisms underlying the expression of ARTs can differ significantly across species. In some cases, tactics are fixed for life (fixed tactics) [6] and often represent distinct life histories. Fixed tactics can occur through either inherited genetic polymorphisms [7–10], condition-dependent switches that are triggered prior to sexual maturation [1,6,11], or a combination of genetic and environmental factors [12,13]. In other cases, individuals can exhibit different tactics throughout their reproductive life, either as they grow or in response to changing social or environmental context (plastic tactics or status-dependent tactics) [1,4,6,14]. Advances in sequencing technology, such as RNA sequencing (RNAseq), now allow behavioral ecologists to explore how variation in gene expression contributes to behavioral variation among mating tactics and examine if the genes influencing these behaviors differ across species with ARTs.

Next-generation sequencing has led to more in-depth research into the molecular mechanisms driving ARTs [9,15–20]. For example, development of independent (territorial) males and two alternative tactics, satellite males and female-mimicking (faeder) males in a shorebird (the ruff, *Philomachus pugnax*) is driven by a supergene resulting from a chromosome inversion that contains 125 predicted genes potentially influencing ART traits [9,10]. However, due to the lack of reference genomes for most teleosts, much of the work on ARTs in this group has focused on examining differential gene expression to identify genes associated with these tactics. Most of these studies have found a large number of genes that vary among tactics in expression in the brain during mating. For example, in the ocellated wrasse (*Symphodus ocellatus*), 1,048 genes were differentially expressed when comparing sneakers to two other male tactics (nesting and satellite) and to females [19]. In the black-faced blenny (*Tripterygion delaisi*) and peacock blenny (*Salaria pavo*), RNAseq identified approximately 600 transcripts differentially expressed within the brains of ‘sneaker’ versus other male tactics [18, 20]. In another study, approximately 2,000 transcripts were differentially expressed between intermediate-sized sailfin molly (*Poecilia latipinna*) males that primarily perform courtship behaviors compared to small males that only perform sneaking behaviors [17]. Changes in social context also led to a larger response (i.e. changes in gene expression) in intermediate-sized males that show higher levels of tactic plasticity when compared to small sneaker males [17], suggesting that genes driving neural response during mating may differ between plastic and fixed tactics.

With the increase in genomic studies examining differences among ARTs, there are a growing number of candidate genes associated with these tactics. Schunter *et al.* [18] proposed a list of potential candidate genes based on a number of studies (Table 1).

**Table 1:** Proposed candidate genes (from [18]) influencing teleost alternative reproductive tactics (ARTs). POA = Pre-optic area

| Proposed Candidate Genes | Function | Relationship to ARTs |
| --- | --- | --- |
| Arginine vasotocin (*avt*) | Non-mammalian homolog of vasopressin. Activates some aspects of sexual behavior | ↑ in posterior POA of territorial cichlid males, but ↑ anterior POA of non-territorial [21]; ↓ density of *avt* mRNA in POA in parental blenny males [22] |
| Gonadotropin releasing hormone (*gnrh*) | Regulates release of luteinizing hormone and follicle-stimulating hormone from the pituitary gland | ↑ in territorial cichlid males [16] |
| Cytochrome P450 family 19, subfamily A, polypeptide 1 (*cyp19a1*) | Brain aromatase. Key enzyme in estrogen biosynthesis | ↑ in territorial cichlid males [16]; ↑ territorial blenny males [23]; ↑ territorial black-faced blenny males [18]; ↓ in the sonic motor nucleus of nesting type I (territorial) male plainfin midshipman compared to type II (female mimic) males [24] |
| Ependymin (*epd*) | Glycoprotein associated with neuroplasticity and neuronal regeneration. Also affects aggression levels in zebrafish [25]; Associated with stress in trout [26] | ↑ in territorial cichlid males [16]; ↓ in subordinate trout males [25] |
| Galanin/GMAP prepropeptide (*gal*) | Neuropeptide that influences neurotransmitters. Associated with male sexual behaviors [27] and parental care [28] | ↑ in territorial cichlid males [16] |
| Somatostatin (*sst*) | Neuropeptide that regulates endocrine pathways. Also affects neurotransmitters | ↑ in territorial blenny males [18]; ↑ in territorial cichlid males [16] |
| Early growth response 1 (*egr1*) | Transcription factor that influences neural plasticity | ↑ when subdominant cichlid males switch to dominant [29] |

This list included gonadotropin releasing hormone (*gnrh*), arginine vasotocin (*avt*), cytochrome P450 family 19 subfamily A polypeptide 1 (*cyp19a1*), ependymin (*epd*), galanin (*gal*), stomatostatin (*sst1* and *sst3*), and early growth response 1 (*egr1*). Many of these genes are known to be involved in hormone regulation and vertebrate mating behavior, and differences in expression levels have been observed among mating tactics in different fish species. For example, the product of the *cyp19a1b* gene is aromatase B, the key enzyme responsible for the conversion of androgens to estrogens within the brain of vertebrates [e.g., 24,30]. *Cyp19a1* also plays an important role in sex determination and sex change in fish [31–33]. Higher levels of *Cyp19a1b* expression have been observed in territorial males compared to sneaker males in the peacock blenny [23], black-faced blenny [18], and an African cichlid (*Astatotilapia burtoni*) [16]. As more data become available, the number of candidate genes in this list will likely increase and evaluating gene expression across teleosts will aid in determining whether similar molecular pathways drive ART behaviors across different species.

One of the best-studied vertebrate species with male ARTs is the bluegill sunfish (*Lepomis macrochirus*). In this species, males have two distinct life histories: parental and cuckolder. In Lake Opinicon (Ontario, Canada), parental males mature at around seven years old and construct nests, court females, and provide care to young [34]. Cuckolder males mature at a significantly younger age, around two years old [34]. Rather than competing with parental males for access to females, cuckolders initially use a “sneaking” tactic to dart in and out of nests while parental males and females are spawning. As they grow, typically around an age of 4 years, cuckolder males appear to transition into “satellite” males by taking on female-like coloration and behaviors [34,35]. Satellite males use this female mimicry to deceive a parental male that he has two true females in his nest [36]. The parental and cuckolder life histories are fixed – once a male adopts the parental or cuckolder life history he remains in that life history [37]. However, within the cuckolder life history, mating tactics are developmentally plastic, with males apparently transitioning from the sneaker tactic to the satellite tactic as they age [37].

While the spawning behavior, reproductive success, and hormone profiles of bluegill have been studied extensively [37–42], the genes influencing behavioral differences during spawning are less clear [43]. Thus, for this study, we used RNAseq to characterize the brain transcriptome of the three spawning male tactics (parental, sneaker, and satellite), non-spawning parental males, and spawning females to examine how differences in gene expression may relate to behavioral variation among these groups. Specifically, we (1) assessed whether or not there is a greater difference in gene expression profiles between fixed tactics (parental versus the two cuckolder tactics) than between tactics within a plastic life history (sneaker versus satellite), (2) identified specific gene categories that are expressed for each tactic, (3) compared expression differences between male and female bluegill, and (4) examined the expression of potential candidate genes identified from other fish species to determine if they also differentiate ARTs in bluegill.

## Materials and Methods

### Bluegill Sampling

In June 2013, bluegill sunfish were collected via dip net from Lake Opinion near Queen’s University Biological Station (QUBS), Elgin, Ontario, Canada. A total of 12 parental males, 12 sneaker males, 13 satellite males, and 12 females were collected directly from the bluegill colony while in the act of spawning. A spawning bout in bluegill typically occurs over the course of one day, following a period of several days of nest construction by parental males. All spawning fish used in this study were behaviorally verified as to tactic before being collected. An additional 12 non-nesting parental males were collected four days prior to spawning (as determined once spawning at these colonies began). Individuals were euthanized using clove oil, total body length was measured, and brains were immediately dissected out and stored in RNAlater (Life Technologies, Carlsbad, CA). Brains remained in RNAlater at 4°C for 24 hours and were then transferred to fresh cryovials, flash fi:Ozen, and kept in liquid nitrogen until they were transported on dry ice to the University of Western Ontario. Samples were then stored at -80°C until RNA extraction. The Animal Care Committee at Western University (UCC) approved all procedures preformed in this study (AUP # 2010-214).

### Total RNA Extraction

Total RNA was extracted using a standard Trizol (Life Technologies, Carlsbad, CA) extraction. RNA was submitted to the London Genomics Center at the University of Western Ontario and quality was assessed using a 2100 Bioanalyzer (Agilent Technologies, Palo Alto, CA). Four individuals from each group (spawning parental males, non-spawning parental males, sneaker males, satellite males, and females), for a total of 20 individuals, were submitted to the Michigan State University Research Technology Support Facility - Genomics Center for cDNA library construction and sequencing. Individuals used for this study had RIN (RNA Integrity Number) values ranging from 9.2-9.9.

### cDNA Library Construction and Sequencing

The cDNA libraries were constructed for each individual using Illumina TrueSeq Stranded mRNA Library Preparation Kits LT (Illumina, San Diego, CA), with each individual receiving a uniquely identifiable index tag. The quality of each library was evaluated and the 20 individuals were multiplexed into a single sample that was subsequently run on two lanes of an Illumina HiSeq2500 Rapid Run flow cell (v1). Sequencing was performed on paired end 2 x 150 bp format reads and bases were called using Illumina Real Time Analysis software (v1.17.21.3). Reads from each individual were identified based on their unique index tag, separated, and converted to fastq files using Illumina Bcl2fastq v1.8.4. Sequencing produced an average of 14.5 million reads per individual, with over 90% of the reads having a Q-score >30.

### *De novo* Transcriptome Assembly and Reference Transcriptome

Prior to assembly, read quality was assessed using FastQC (http://www.bioinformatics.babraham.ac.uk/projects/fastqc). Nucleotides whose quality score was below PHRED=2 were trimmed using Trimmomatic version 0.32 [44], following recommendations from MacManes [45]. The reference transcriptome was assembled *de novo* using Trinity version 2.04 [46,47]. One representative of each of the five groups (spawning parental male, non-spawning parental male, sneaker male, satellite male, and female) was used to construct a combined reference transcriptome. The five representatives selected for the reference were the individuals with the highest number of reads within their group and, a total of 85 million paired-end reads were assembled. The assembly was conducted with both normalized and non-normalized reads and normalization was performed using Trinity’s *in silico* normalization program. To test the completeness of the transcriptome, reads from samples not used in the assembly were mapped back to the transcriptome using Burrows-Wheeler Aligner (bwa)-mem version 0.7.12 [48], and >90% of those reads aligned, which is comparable to the rate of mapping for the individuals used in the assembly (92%).

TransDecoder [46] was used to identify protein-coding regions within the assembled transcriptome. Transcripts that contained protein-coding regions or transcripts that blasted to compete coding sequences (cds) and non-coding RNA (ncRNA) from spotted green puffer (*Tetraodon nigroviridis*), spotted gar (*Lepisosteus oculatus*), southern platyfish (*Xiphophorus maculatus*), medaka (*Oryzias latipes*), Japanese pufferfish (*Takifugu rubripes*), West Indian Ocean coelacanth (*Latimeria chalumnae*), Mexican tetra (*Astyanax mexicanus*), zebrafish (*Danio rerio*), or Amazon molly (*Poecilia formosa*) (downloaded from Ensembl) comprised the reference transcriptome used for both read alignment and to estimate transcript counts.

### Read Alignment and Transcript Counts

Reads from each individual were separately aligned to the reference transcriptome using bwa-mem 0.7.10 [48]. At least 85% of all reads from each individual mapped back to the reference, with the majority aligning 90% of reads or higher. The sequence alignment/map (sam) files were then converted to a binary format (bam) using Samtools 0.1.19 [49]. Transcript counts for each individual were obtained using the program eXpress 1.5.1 [50]. Differential gene expression was determined using the R statistical package edgeR 3.6.8 [51]. Low abundance transcripts were filtered out, leaving 19,804 transcripts for differential analysis. Transcript counts were normalized to account for differences in cDNA library size among individuals and dispersion parameters were estimated using Tagwise dispersion estimates. Differences in gene expression comparing paired treatments were calculated using an Exact-test for binomial distribution. Genes with p-values lower than 0.05 after false discovery rate (FDR) correction were determined to be statistically significant. All fold changes are reported as log2 fold change. Genes with FDR values below 0.05 and with log2 fold changes greater than 1.5 were used for hierarchical cluster analysis to examine overall group differences.

### Gene Annotation and Enrichment Analysis

Both the reference transcriptome and transcripts differentially expressed among groups were blasted using Blastx against a custom-assembled fish protein database. This database consisted of Ensembl protein databases of 13 different fish species: Amazon molly (*Poecilia formosa*), zebrafish (*Danio rerio*), Mexican tetra (*Astyanax mexicanus*), Atlantic cod (*Gadus morhud*). West Indian Ocean coelancanth (*Latimeria chalumnae*), Japanese pufferfish (*Takifugu rubripes*), sea lamprey (*Petromyzon marinus*), medaka (*Oiyzias latipes*), southern platyfish (*Xiphophorus maculatus*), spotted gar (*Lepisosteus oculatus*), three-spined stickleback (*Gasterosteus aculeatus*), green spotted pufferfish (*Tetradon nigroviridis*), and Nile tilapia (*Oreochromis niloticus*). Blast hits with e-values less than 1x10^−10^ were considered significant. Ensembl IDs from the blast hits were then converted into GO term identifiers using Biology Database Network (bioDBnet) (http://biodbnet.abcc.ncifcrf.gov/db/dbFind.php).

For purposes of gene annotation and enrichment analysis, we focused on transcripts within the reference transcriptome that were not filtered out of the data set due to low transcript expression (total of 19,804 transcripts). To examine which GO terms were significantly enriched within this set, unique Ensembl IDs from Blastx were converted to Ensemble IDs associated with stickleback homologs using the R package biomaRt 2.20.0. Enrichment analysis relative to the stickleback genome was then conducted on these homologs using the BioMart portal (http://central.biomart.org/enrichment).

For the transcripts that were differentially expressed among behavioral groups, enrichment analysis was conducted using a Fisher Exact test to examine whether the proportion of genes within each GO category was significantly higher than what would be expected based upon the proportion of genes assigned to that GO term within the reference transcriptome. To ensure adequate statistical power, only GO terms with at least 10 transcripts within each category were included in the statistical analysis. A FDR correction was applied to control for multiple testing and GO terms with p-values < 0.05 were considered to be significant. Visual representations of enriched GO terms were generated using REVIGO [52].

## Results

### Reference Transcriptome

This study presents the first reference transcriptome for the brain of bluegill sunfish. The fully assembled transcriptome consisted of 272,189 transcripts. Of these, 72,189 transcripts contained cds or blasted to ncRNA from the customized Ensembl fish database. These 72,189 transcripts were then used as the reference transcriptome for alignment and mapping. The mean transcript length within the reference transcriptome was 2,024 bp, with N50 = 3,106 bp and N90 = 1,018 bp. The largest transcript consisted of 27,880 bp. Approximately 82% of the transcripts had only one isoform, while 18% (12,951 transcripts) had two or more isoforms.

For GO enrichment analysis, we only examined the 19,804 transcripts within the reference transcriptome that passed our established filtering process. Of these, 18,104 had significant Blastx hits with Ensembl gene IDs (S1 Table), of which 12,224 transcripts had stickleback homologs that could be used to examine GO term enrichment for the bluegill brain transcriptome compared to the stickleback genome. The GO terms with significant enrichment for the bluegill reference transcriptome included translation, catabolism, vesicle-mediated transport, biosynthesis, small molecule metabolism, and generation of precursor metabolites and energy (Figure 1A & B).

**Fig. 1:**
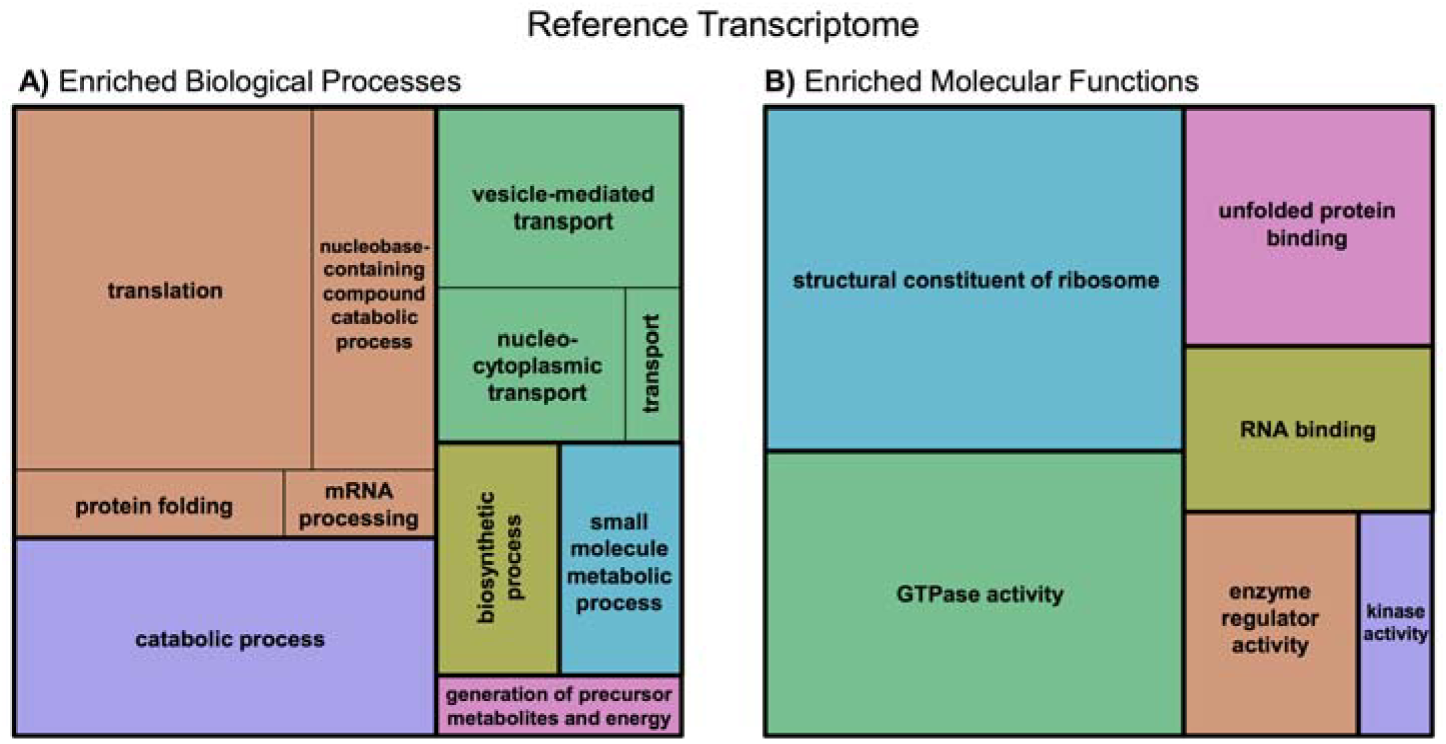
Significant GO terms associated with the reference transcriptome. GO terms related to (A) biological processes and (B) molecular functions that were significantly enriched in the bluegill reference transcriptome relative to the stickleback genome. Boxes of similar color can be grouped into the same GO term hierarchy. The size of each box reflects the - log10 p-value of the GO term within each group.

### Differential Gene Expression across All Groups

Based on hierarchical cluster analysis, sneaker males grouped separately from the other male tactics (Figure 2). Spawning parental males, non-spawning parental males, and females displayed similar expression profiles to each other. Satellite males tended to have expression profiles intermediate of sneakers and the other groups.

**Fig. 2:**
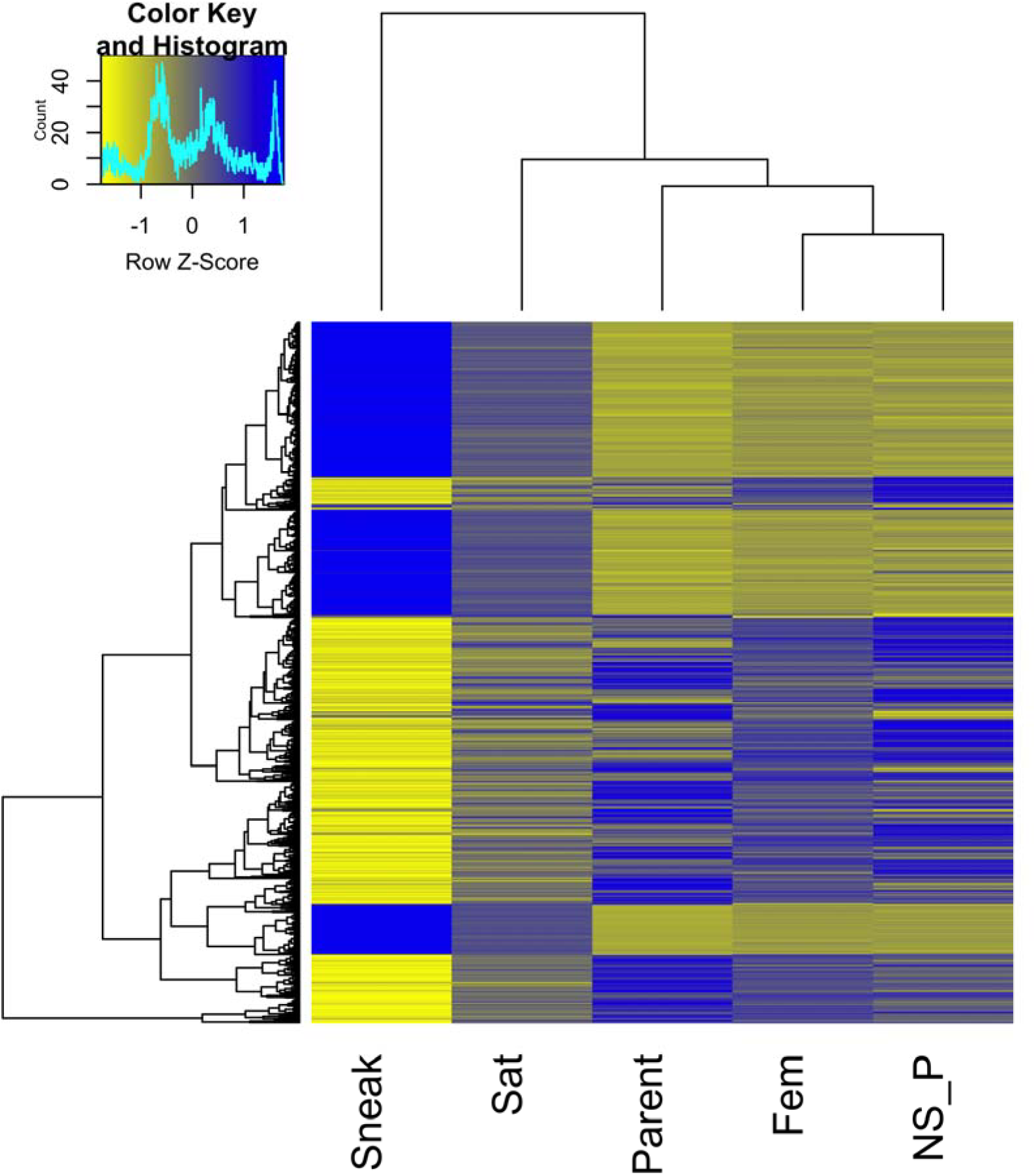
Heatmap of transcripts differentially expressed between at least one group comparison. Only transcripts with a log2 fold change of 1.5 or greater are included and 1,400 transcripts are represented. Expression values are scaled by row. Sneak = sneaker males, Sat = satellite males. Parent = parental males, Fem = females, NS_P = non-spawning parental males.

When comparing across all groups, five unique transcripts consistently displayed higher expression in spawning parental males compared to all other groups (Table 2). Fourteen unique transcripts were differentially expressed in satellite males compared to all other groups. Expression for these transcripts in satellite males was higher compared to parental males (spawning and non-spawning) and females, but lower compared to sneaker males (Table 2). There were 2,253 transcripts differentially expressed between sneaker males and all other groups (S2 Table). The majority of these transcripts with higher expression in sneakers were related to ion transport, ionotropic glutamate signaling pathway, and mRNA processing (Figure 3). Two transcripts were differentially expressed in females compared to the other groups and both of these were expressed at lower levels than in the other groups (Table 2).

**Table 2:**
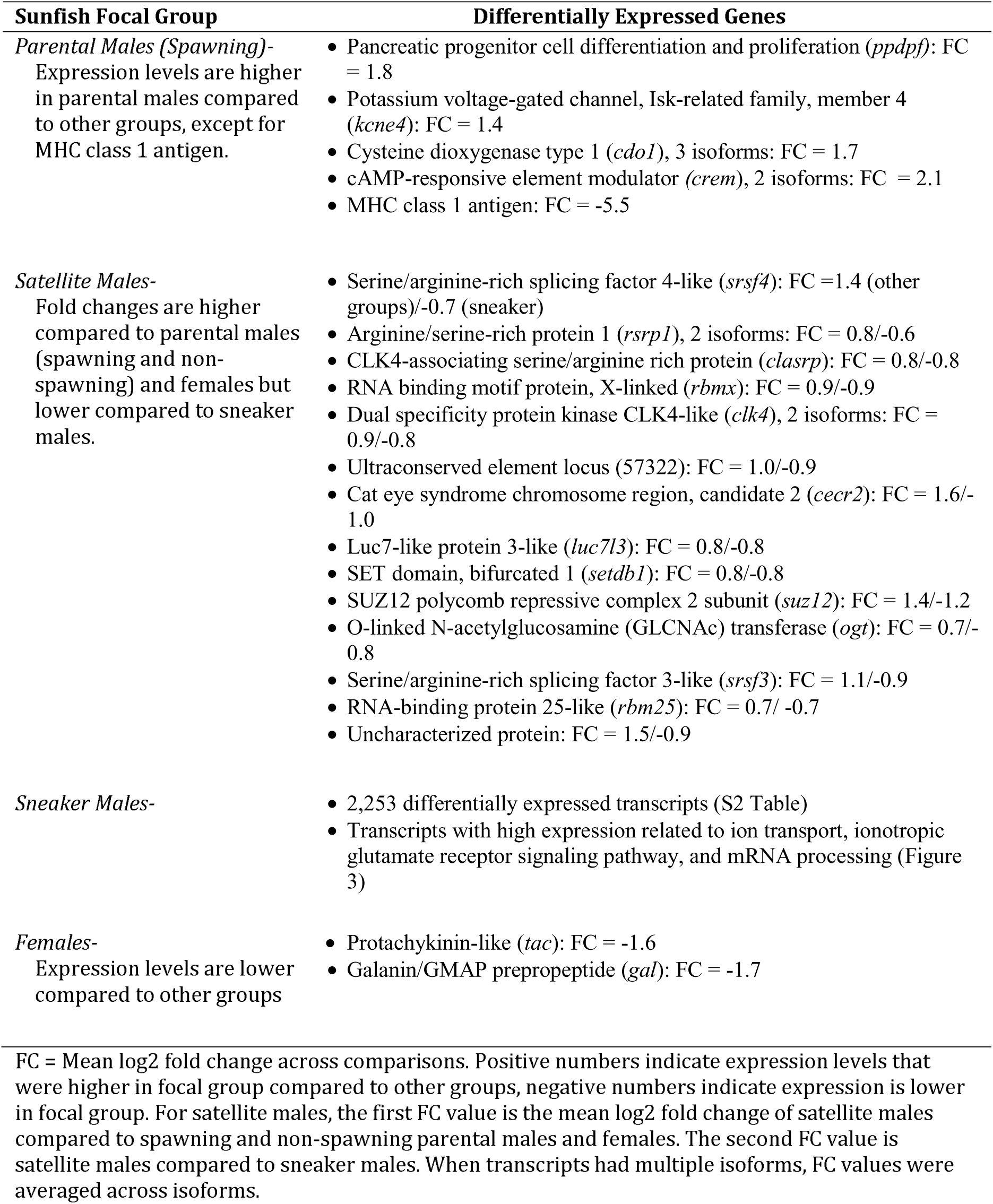
Differentially expressed transcripts associated with each male mating tactic and females.

**Figure 3:**
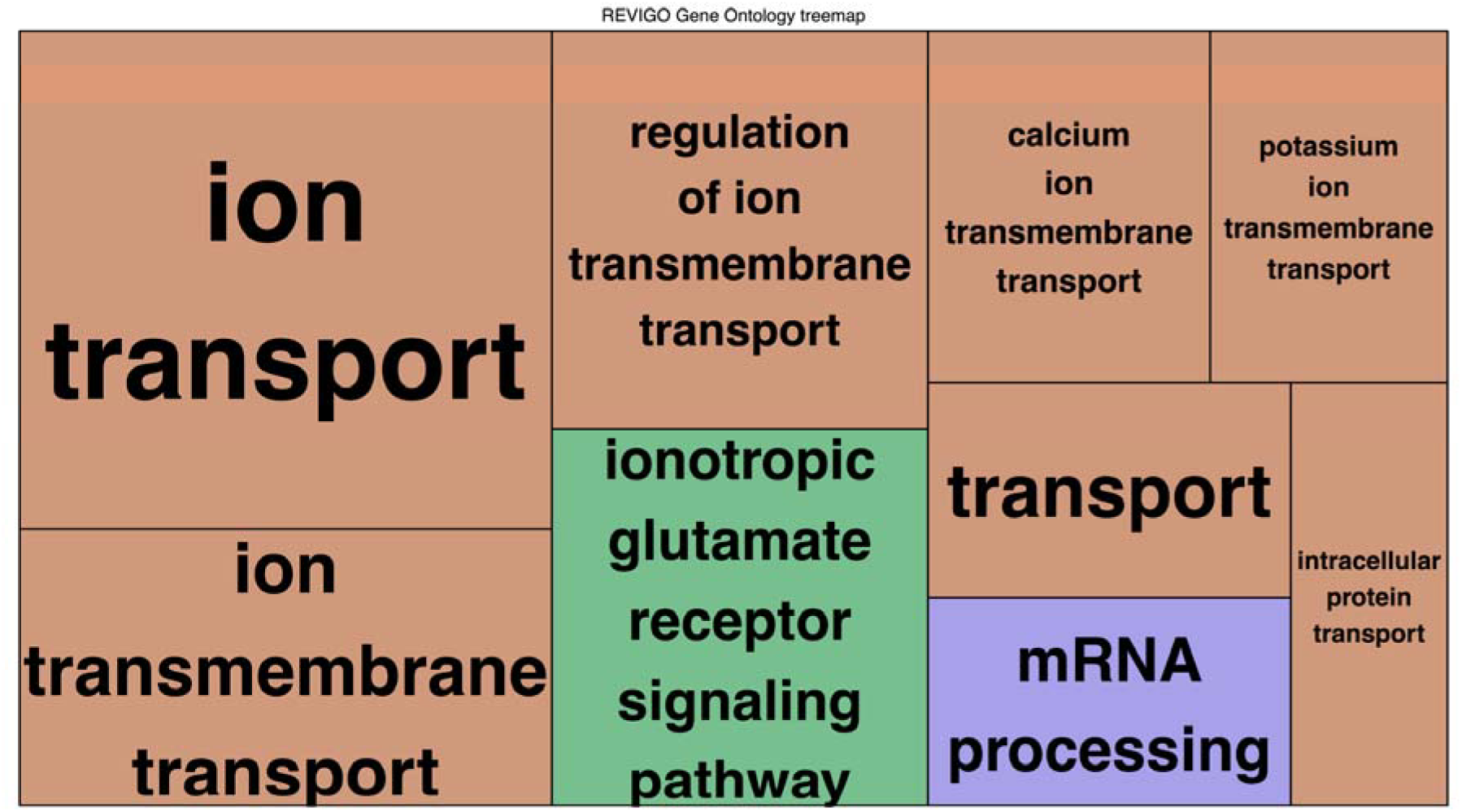
Biological process GO terms enriched by genes with higher expression in sneaker males compared to all other groups. Boxes of similar color are grouped into the same GO term hierarchy. Box size reflects the - log10 p-value of the GO term.

### Between Life History Comparisons

*Spawning Parental Males versus Sneaker Males*. A total of 9,279 transcripts were differentially expressed between spawning parental males and sneaker males. Of these, 4,537 transcripts showed higher expression in parental males (S3 Table) and 4,742 transcripts showed higher expression in sneaker males (S4 Table).

Enrichment analysis of GO terms associated with differentially expressed genes showed that the biological functions most enriched in parental males included translation, translational initiation, translation elongation, proteolysis involved in cellular protein catabolism, and oxidation-reduction processes (S5 Table). The 27 molecular processes most enriched in parental males compared to sneaker males included ribosomal structure, oxidoreductase activity and catalytic activity (S5 Table).

Biological processes enriched with genes displaying higher expression in sneaker males included ion transport, homophilic cell adhesion, protein phosphorylation, ionotropic glutamate receptor signaling pathway, and synaptic transmission (S5 Table). The 10 molecular processes enriched in sneaker males included ion channel activity, protein binding, ionotropic glutamate receptor activity, and extracellular glutamate-gated ion channel activity (S5 Table).

*Spawning Parental Males versus Satellite Males*. A total of 1,141 transcripts were differentially expressed between spawning parental males and satellite males. Of these, 676 transcripts had higher expression in parental males (S6 Table) and 465 transcripts showed higher expression in satellite males (S7 Table).

Only one GO term related to biological function, oxidation-reduction processes, was enriched in parental males compared to satellite males (S5 Table). Six GO terms related to molecular processes were enriched in parental males relative to satellite males (S5 Table). These were iron ion binding, two types of oxidoreductase activity, heme binding, acylCoA dehydrogenase activity and catalytic activity (S5 Table).

Only one GO term related to biological function, ion transport, was enriched in satellite males compared to spawning parental males (S5 Table). Three GO terms related to molecular processes were enriched in satellite males relative to spawning parental males. These were nucleic acid binding, ion channel activity, and GTP binding (S5 Table).

### Differential Expression within Life Histories

*Satellite Males verses Sneaker Males*. There were 2,590 transcripts differentially expressed between satellite males and sneaker males. Of these, 2,480 transcripts were also differentially expressed between spawning parental and sneaker males and all showed expression to be in the same direction for parental and satellite males compared to sneakers (i.e. those with higher expression in parental males compared to sneaker males were also higher in satellite males compared to sneakers). Only 110 transcripts were differentially expressed in satellite males compared to sneaker males that were not also differentially expressed between parental and sneaker males. Seventy-six transcripts had higher expression levels in satellite males (S8 Table) and 34 transcripts had higher expression in sneaker males (S9 Table). The number of transcripts differentially expressed was too low to have adequate statistical power to perform enrichment analysis for GO terms. However, many of the transcripts with higher expression in satellite males are associated with GTP catabolism, while many transcripts with higher expression in sneaker males are involved in signal transduction, neural crest cell migration, and DNA integration.

*Spawning Parental Males verses Non-Spawning Parental Males*. A total of 137 transcripts were differentially expressed between spawning and non-spawning parental males. The majority of these transcripts (132 transcripts) showed higher expression in spawning males (S10 Table). Genes with the highest expression in spawning parental males compared to non-spawning males were MHC II beta antigen, cytosolic 5′-nucleotidase II (*nt5c2*), cAMP responsive element modulator a (*crem*), cysteine dioxygenase type 1 (*cdo1*), and an uncharacterized protein. Only 8 transcripts showed higher expression in non-spawning parental males. These were nuclear receptor subfamily 1 group D member 4b (*nr1d4b*), neuronal tyrosine-phosphoinositide-3-kinase adaptor 2 (*nyap2*), sphingosine-1-phosphate receptor 4 (*s1pr4*), gamma-aminobutyric acid A receptor beta 3 (*gabrb3*), and four uncharacterized proteins (S11 Table). Again, the number of transcripts assigned to GO term was too small to have adequate statistical power to perform an enrichment analysis for this comparison.

### Sex Differences

Two unique transcripts were differentially expressed between females and all of the male groups (sneaker, satellite, spawning parental male, and non-spawning parental males) (Table 2). These corresponded to galanin/GMAP prepropeptide (*gal*) and protachykinin (*tac*) and both were expressed at lower levels in females. Expression profiles of females were more similar to spawning and non-spawning parental males than to satellite males, despite females’ and satellites’ similarity in spawning behavior (Figure 2).

### Potential Candidate Genes Associated with ART Spawning Behavior

We observed differential expression of a number of transcripts previously identified as potential candidate genes associated with differences in ART spawning behaviors (described in Table 1) (Table 3). In our data set, the candidate genes *Cyp19a1b, epd*, and *gal* showed higher expression in spawning parental males compared to sneaker males. *Epd* also had higher expression in satellite males compared to sneakers. *Egr1* showed higher expression in both satellite and sneaker males relative to spawning parental males. *Sst1* showed higher expression in satellite males compared to sneaker males, but no differences in other comparisons between tactics. No differences in expression related to *gnrh, avt*, or *sst3* were observed between any of our groups.

**Table 3:** Gene expression differences (log2 fold change) among male tactics for proposed candidate genes (see Table 1).

| Proposed Candidate Gene | Isoform ID | Comparison between Male Tactics (Log2 Fold Change) |  |  |  |
| --- | --- | --- | --- | --- | --- |
|  |  | Parent vs Sneak | Parent vs Sat | Sat vs Sneak | Spawn Parent vs NonSpawn |
| Arginine vasotocin (*avt*) | c34708_g2_i1 | 0.45 [0.32] | -0.98 [0.09] | 0.54 [0.33] | 0.74 [0.50] |
| Gonadotropin releasing hormone (*gnrh*) | c63124_g1_i1 | 0.76 [0.50] | 0.32 [0.87] | 0.44 [0.77] | 0.77 [0.88] |
| Cytochrome P450 19a1b (*cyp19a1b*) | c48084_g2_i1 | **0.93 [0.0002]** | 0.64 [0.06] | 0.28 [0.40] | 0.39 [0.58] |
| Ependymin (*epd*) | c44195_g1_i5 | **1.54 [1.4 x 10<sup>-8</sup>]** | 0.66 [0.07] | **0.89 [0.007]** | -0.51 [0.45] |
| Galanin/GMAP prepropeptide (*gal*) | c41071_g5_i2 | **1.12 [0.0001]** | 0.53 [0.91] | -0.59 [0.10] | 0.09 [0.97] |
| Somatostatin 1 (*sst1*) | c30013_g1_i1 | 0.53 [0.15] | -0.39 [0.49] | **0.93 [0.03]** | 0.27 [0.88] |
| Somatostatin 3 (*sst3*) | c46547_g6_i1 | 0.001 [1.00] | -0.25 [0.54] | 0.25 [0.48] | 0.15 [0.90] |
| Early growth response 1 (*egr1*) | c37907_g1_i1 | **-0.74 [0.02]** | **-0.91 [0.03]** | 0.16 [0.72] | -0.63 [0.42] |

Values in brackets represent p-values after false discovery rate correction. Values in bold are significant at $p < 0.05$ . Parent = parental male, Sneak = sneaker male, Sat = satellite male, NonSpawn = non-spawning parental male.

In addition to these previously identified candidate genes, another transcript that displayed large differences in expression between spawning parental males and all other groups (including non-spawning males) was associated with cAMP-responsive element modulator (*crem*) (Table 2). Multiple isoforms of the transcript were expressed, with log2 fold changes ranging from 1.3 - 2.6 times higher in spawning parental males compared to other groups. Consistent with the finding for GO term enrichment, transcripts that showed the highest levels of expression in sneaker males compared to other groups were related to glutamate receptor genes, particularly AMPA ionotropic glutamate receptors (S2 Table).

In addition to the candidate genes listed in Table 1, a number of endocrine genes were differentially expressed among two or more male tactics. Among these genes are a number of genes that we consider candidate genes based on documented male tactic differences in circulating steroid hormone levels on the day of spawning [38,39,42]. Further investigation of these genes is currently in progress and will be reported elsewhere.

The datasets supporting the conclusions of this article are available on the Sequence Read Archive (SRA) through BioProject ID: PRJNA287763. Environmental data, RNA quality information, the assembled transcriptome, the transcript count matrix, and R code for differential gene analysis are currently available for review at the following link (https://www.dropbox.com/sh/jxbgsiyrz89npfn/AAA5n-iNR0zhHcPcEPrlCT72a?dl=0) and will be available on Dryad upon acceptance.

## Discussion

Bluegill sunfish are a classic system for examining behavioral differences in ARTs. In this study, we generated and assembled the first bluegill brain transcriptome and identified candidate genes that contribute to differences in male spawning tactics. The main differences in gene expression were found between sneaker males when compared to the two other male tactics and females. Generally, sneaker males showed higher expression in transcripts influencing neural activity, whereas parental and satellite males exhibited higher expression in genes related to translation and oxidoreductase activity. There were larger differences in transcript expression among different male tactics than between males and females.

### Overall Expression Differences among ARTs

One of our main findings is that a shared life history does not appear to be a driving factor influencing similarity in neural gene expression among male tactics.

In bluegill, parental and cuckolder life histories are fixed, but within the cuckolder life history, males transition from the sneaking to the satellite tactic as they age [34,38]. Our data showed that, regardless of whether comparisons were made across fixed (parental versus sneaker or parental versus satellite) or plastic (sneaker versus satellite) tactics, sneaker males showed the highest level of differentiation in gene transcription. Similar results have been observed in the ocellated wrasse, where sneaking males also showed greater differences compared to other tactics [19]. The expression differences in sneakers may be partially due to age because sneaker males in both bluegill and ocellated wrasse are typically younger than satellite and parental or territorial males (although this is not always the case for *S. ocellatus* [53]). Genes associated with increased age in other fish species, such as translation elongation and ribosomal proteins [54], had higher levels of expression in parental and satellite males compared to sneaker males in our dataset. In addition to age, differences in expression may also be characteristic of behavioral differences. Behaviors exhibited by sneaker males during spawning usually differ in fundamental aspects from those of other male tactics. In the ocellated wrasse, for example, satellite and nesting males cooperatively protect the nest from sneakers and other egg predators [55], potentially leading to similar expression profiles between these tactics. In bluegill, satellite and parental males associate closely with the female throughout spawning, whereas sneakers dart in and out of the nest. These differences in spawning tactics might also contribute to the differences in gene expression observed in the two studies. Thus, age and spawning tactic are likely important contributors to gene expression patterns across ARTs, and life history is not exclusively responsible for these differences.

### Gene Categories Associated with ARTs

Identifying distinct gene categories expressed by ART types provides information regarding which gene classes influence behavioral differences during spawning. Previous studies in sailfin mollies, *Poecilia latipinna*, and Atlantic salmon. *Salmo salar*, indicate that sneaker males have increased expression of genes related to neurotransmission and learning [15,17]. We found that the GO terms enriched in bluegill sneaker males compared to all other groups were the ionotropic glutamate signaling pathway and ionotropic glutamate receptor activity. Ionotropic glutamate receptors are primarily excitatory neurotransmitter receptors and play an important role in fast synaptic transmission [reviewed in 56]. Two of these receptors, NMDA and AMPA, play important roles in memory function and spatial learning [reviewed in 57]. Blocking NMDA receptors impairs learning new spatial locations in goldfish [58] and mice with impaired AMPA receptors show normal spatial learning but have impaired working spatial memory (i.e. their ability to alter their spatial choice in response to changing environments is impaired) [59]. We propose that increased expression of genes related to spatial memory, particularly working spatial memory, could be important for bluegill sneakers during spawning as they attempt to gain access to nests while avoiding detection not only by the parental males, but also common predators around the colony [60]. Bluegill sneakers must also position themselves in close proximity to females so they can time sperm release to coincide with female egg release [61]. Similarly, sailfin molly sneakers, who also show enrichment in ionotropic glutamate related genes [17], probably benefit from increased working spatial memory as they position themselves by the female for quick and successful copulations. In this context, increased expression in gene pathways that improve neural function related to working spatial memory would be especially beneficial for sneaking tactics to increase their reproductive success.

While ARTs with fixed tactics maintain the same mating tactic over their lifetime, ARTs with plastic tactics can alter their behavior and, in some cases their phenotype, when switching from one tactic to another. Diverse phenotypes can be accomplished without altering the underlying genomic sequence through a number of mechanisms including epigenetic regulation, alternative gene splicing, and post-translational modification of proteins. A number of genes involved in these processes showed higher expression in the plastic tactics (satellite and sneaker) compared to the fixed parental tactic (Table 2). For example, *ogt* plays a key role in chromatin restructuring and post-translational modification of proteins [62]. It has been also implicated in a number of different processes including nutrient and insulin signaling [63,64], sex-specific prenatal stress [65], and behavioral plasticity [66]. Genes associated with alternative splicing that were expressed at higher levels in plastic tactics included isoforms of serine/arginine-rich proteins (SR proteins), a family of proteins involved in RNA splicing [67], and CLK-4 like proteins, which are kinases that function in regulating SR protein activity [68]. Similarly, differential expression of RNA splicing genes have also been observed in two other teleost species with plastic tactics, the black-faced blenny and intermediate-sized sailfin mollies [17,18]. While the mechanisms influencing how ART males switch between tactics is currently unresolved, epigenetic regulation, alternative gene splicing, and post-transcriptional modifications could be important for plastic tactics in altering their phenotype in response to environmental or developmental cues.

### Sex Differences

Neural differences between the sexes are common and found in many taxa [reviewed in 69,70]. However, within ARTs, differences in neural expression profiles can often be larger among male tactics than between males and females [18–20]. In bluegill, only two transcripts were differentially expressed in females compared to all male groups and these corresponded to *gal* and *tac*. *Gal* and *tac* are neuropeptides and neurons expressing these genes have been associated with male sexual behavior and aggression [27, 71]. Injections of *gal* into the preoptic area (MPOA) of the brain increase sexual behaviors in male rats [27] and stimulate both male-typical and female-typical sexual behaviors in females [72]. In male rats, testosterone can enhance the pituitary’s response to *gal*, which heightens *gnrh's* stimulation of luteinizing hormone (LH). If *gal* is directly involved in regulating *gnrh* response in bluegill, this neuropeptide may play an important role in behavioral differences between the sexes. In sequential hermaphroditic fish, surges in *gnrh* drive the switch from female to male [73]. While bluegill are gonochoristic, gonadal sex is not evident until 30-60 days post hatch [74] and changes in sex can be hormonally induced [75]. Thus, *gal* expression, through its influence on *gnrh*, may play an important role in sex differences for this species. In addition, *gal* also stimulates feeding in fish [reviewed in 76] and feeding may indeed differ between female and male sunfish in the days leading up to and including spawning.

The role of *tac* in influencing sexual behaviors in teleosts has not been addressed, but *tac* expression significantly increases in the brain of male eels during sexual maturation [77] and leads to increased male aggression in *Drosophila* [71]. In bluegill, the primary role of *tac* may not be male-male aggression considering higher expression levels of this gene are also observed in the non-aggressive sateUite and sneaker males when compared to females. Although the ways in which *gal* and *tac* specifically influence sex-specific behaviors in bluegill is currently undefined, the fact that lower expression is consistently observed in females compared to all male groups suggests they are important sex-specific neural genes.

### Candidate Genes Associated with ARTs

A number of candidate genes have been proposed to influence the expression of ARTs in teleosts [18] (Table 1). In our study of bluegill, we corroborate some of these candidates. For example, *Cyp19a1b, epd*, and *gal* had higher expression levels in spawning parental males compared to sneaker males. The expression patterns for all three genes are similar to what has been observed in cichlids [16]. In addition, expression of *epd* is lower in rainbow trout, *Oncorhynchus mykiss*, males that use a sneaking tactic versus males that are dominant and territorial [25], which is also consistent with our findings. In contrast, the one candidate gene that was expressed opposite to expectations was *egr1*. *Egr1* expression was lower in bluegill spawning parental maies compared to sneaker or satellite males although previous work in cichlids found that expression of this gene increases when subdominant males transition into dominant males [29]. *Egr1* is an important transcription factor involved in neural plasticity [78], so it may be one of a group of genes involved in regulating the switch from one tactic to another. Taken together, our results corroborate roles for *Cyp19a1b, epd, gal*, and *egr1* as candidate genes contributing to behavioral differences in ARTs across species. This study is a first step in examining how bluegill ARTs differ in neural gene expression during spawning, and future work will explore how candidate genes are expressed across different brain regions, as some studies have found regional differences associated with genes, such as *avt*, in other species with ARTs [21, 79–82].

We also identified one transcript with a previously unrecognized function in influencing male spawning behavior for any teleost. Transcripts corresponding to isoforms of *crem* were expressed at significantly higher levels in spawning parental males compared to all other male groups, including non-spawning parental males. *Crem* plays a key role in modulating the hypothalamic-pituitary-gonadal axis by regulating transcriptional responses to cAMP in neuroendocrine cells and also serves as an important activator of spermatogenesis in Sertoli cells of mice [83–85]. This gene can act as both transcriptional activator and inhibitor depending on the splice variant produced [83]. One splice variant is inducible cAMP early repressor (ICER), a powerful repressor of cAMP-regulated transcription [86]. ICER plays a key role in circadian melatonin synthesis by repressing the key enzyme that converts serotonin to melatonin [87]. Both serotonin and melatonin can influence behavioral responses in fish [88–91]. High levels of these neurotransmitters have been associated with increased mating and cooperative behavior and decreased aggressive behavior [88,90,91]. ICER has not yet been well characterized in teleosts but one of our differentially expressed *crem* transcripts had a significant blast hit to an ICER variant from *Epinephelus brunes* (longtooth grouper). The relationship among *crem*, melatonin, and aggression is opposite to what would be expected if ICER is playing a role since parental males have darker pigmentation and are more aggressive than other groups [60, 92–94]. However, increased expression of *crem*, whether through ICER or another *crem* transcript variant, could be a candidate gene influencing behaviors associated with parental male spawning given its role in transcriptional regulation and its involvement in the hypothalamic-pituitary-gonadal axis.

In summary, our work describes differences in gene expression profiles in the brains of bluegill sunfish during spawning. The largest differences in expression levels were observed when comparing sneakers to parental males, satellite males, and females, suggesting that differences in gene expression are more related to male reproductive tactic than to life history. Consistent with other studies, our work demonstrates that sneaker males have greater expression of genes involved in neural function relative to more territorial-type males, particularly in relation to working spatial memory, as mediated by ionotropic glutamate receptors. We also found support for the previously identified candidate genes *Cyp19a1b, epd, gal*, and *egr1* contributing to behavioral differences in ARTs and identified a potential new candidate gene, *crem*, for regulating parental males’ behavior during spawning.

## Acknowledgments

We thank Scott Colborne for his help in collecting bluegill, Dave Bridges for providing the R script to convert Ensemble IDs to stickleback homologs, and David Winter and Jeramia Ory for providing Python script used in the bioinformatics analyses. We also thank Shawn Garner, Tim Hain, Lauren Kordonowy, and Lindsay Havens for helpful comments on the manuscript.

## Supporting files

S1 Table: Annotated reference transcriptome

S2 Table: Transcripts differentially expressed in sneaker males compared to all other groups.

S3 Table: Transcripts with significantly higher expression in bluegill parental males compared to sneaker males.

S4 Table: Transcripts with significantly higher expression in bluegill sneaker males compared to parental males.

S5 Table: Biological process and molecular function GO terms that are significantly enriched with genes differentially expressed between tactics.

S6 Table: Transcripts with significantly higher expression in bluegill parental males compared to satellite males.

S7 Table: Transcripts with significantly higher expression in bluegill satellite males compared to parental males.

S8 Table: Transcripts with significantly higher expression in bluegill satellite males compared to sneaker males.

S9 Table: Transcripts with significantly higher expression in bluegill sneaker males compared to satellite males.

S10 Table: Transcripts with significantly higher expression in spawning parental males compared to non-spawning parental males.

S11 Table: Transcripts with significantly higher expression in non-spawning parental males compared to spawning parental males.

